# *Pml* loss worsens NEK1-linked ALS and *Pml* induction drives NEK1 degradation, precluding disease onset

**DOI:** 10.1101/2024.11.23.622051

**Authors:** Panagiota Georgiadou, Bahriye Erkaya, Michiko Niwa-Kawakita, Merve Oltan, Yigit Kemal Keskin, Egemen Sahin, Harun Öztürk, Fatmanur Tiryaki, Kutay Yildiz, Idil Özgenç, Emre Pekbilir, Sukru Anil Dogan, Valérie Lallemand-Breitenbach, Stephanie Vargas, Alain Prochiantz, Elif Nur Firat-Karalar, Hugues de Thé, Umut Sahin

## Abstract

Germinal mono-allelic loss-of-function mutations of *NEK1* drive Amyotrophic Lateral Sclerosis (ALS) at variable penetrance, presumably through haploinsufficiency. Modeling the ALS-associated Arg812Ter mutation in mice revealed that the resulting truncated Nek1 (Nek1^t^) is aggregation-prone, particularly in alpha-motoneurons (αMNs), and drives canonical ALS symptoms when bi-allelically expressed (*Nek1^t/t^)*. Promyelocytic leukemia (*Pml*) ablation allows for ALS symptoms to occur even in heterozygote *Nek1^wt/t^* animals, mimicking the human situation. *Pml* precludes disease occurrence by promoting SUMO-facilitated degradation of Nek1^t^ proteins through PML nuclear bodies (NBs). Conversely, *Pml* induction, achieved by activating the interferon pathway via poly(I:C) treatment, clears Nek1^t^ aggregates in αMNs, dramatically reducing ALS-associated symptoms and extending survival by 5 months. Our studies highlight the role of NEK1 aggregates in ALS pathogenesis and identifies activation of interferon pathways as a candidate therapeutic strategy that not only promotes *Pml-*triggered SUMOylation/degradation of toxic misfolded proteins *in vivo*, but also facilitates the clearance of protein aggregates, yielding dramatic clinical improvement. These observations validate PML as a relevant therapeutic target in neurodegenerative conditions associated with protein aggregation.

## INTRODUCTION

Amyotrophic Lateral Sclerosis (ALS), the third most common neurodegenerative disorder in humans, is a neuromuscular pathology, usually fatal 3-4 years after symptoms onset due to brainstem and spinal alpha-motoneuron (αMN) retrograde degeneration ^1^. Presently, treatment options for ALS are limited and lack effectiveness.

Approximately 5-10% of ALS cases exhibit familial traits, characterized by Mendelian inheritance patterns. The other 90-95% is sporadic with no family history ^2^. The genetic landscape of ALS is complex; for instance, familial ALS forms may be caused by mutations in over 30 genes including *SOD1*, *C9orf72*, *FUS* and *TDP43BP* ^3^. Understanding the cellular and molecular mechanisms driving disease pathogenesis has been challenging due to the diverse functional and structural nature of the proteins encoded by these genes.

A novel ALS-associated gene, *NEK1,* encodes a dual serine-tyrosine kinase (NIMA-related kinase 1 or NEK1), which, predominantly resides in the cytoplasm and associates with the primary cilium ^4^. Following DNA double-strand breaks, NEK1 translocates to the nucleus, co-localizes with DNA damage markers, and activates checkpoint kinases CHK1 and CHK2 for proper DNA damage response ^5,6^. Aside from polycystic kidney disease in mice, NEK1 is linked to other ciliopathies *in vivo*, including short-rib thoracic dysplasia and oral-facial-digital syndrome type II in humans ^7–11^. *NEK1* has emerged as an important ALS-associated gene, with mono-allelic loss-of-function variants in 2-5% of familial, but also sporadic forms ^3,4,12–15^. Recent studies have highlighted the contribution of defective NEK1 activity in ALS development ^12^. In particular, several ALS-associated *NEK1* mutations introduce premature stop codons, such as Arg812Ter, potentially encoding truncated dysfunctional NEK1 variants (NEK1^t^) (Figure 1A) ^13,16^. In principle, these NEK1 variants could also exert dominant-negative effects, gain of function or yield haploinsufficiency. As in other neurodegenerative diseases, protein aggregation is a common phenomenon in ALS ^17^. Nevertheless, whether misfolding-induced aggregation is implicated in mutant *NEK1*-driven ALS remains unclear ^3,4,13,14,16^.

**Figure 1:**
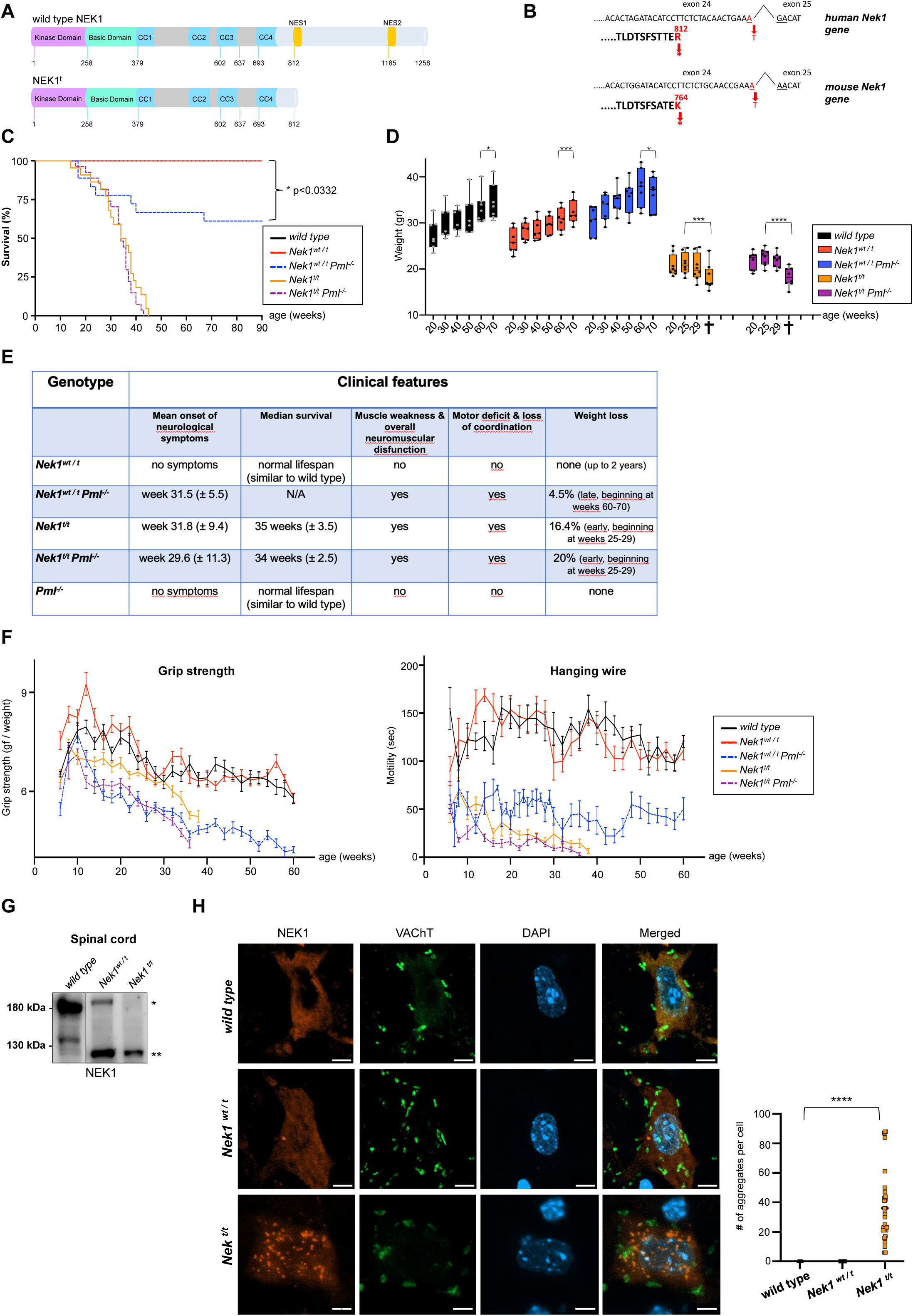
*Pml* safeguards against development of ALS driven by a truncated NEK1 mutant aggregating in αMNs. **A)** NEK1^t^ domain structure compared with wild-type NEK1. CC: coiled coil. NES: nuclear export sequence. **B)** Genomic location of the point mutation introduced in the mouse *Nek1* gene, compared to ALS patients. Amino acids in bold. **C)** Survival of *Pml*-proficient or -deficient *Nek1^wt/t^* or *Nek1^t/t^* mice. For clarity, *Pml^-/-^* controls are presented in Supplementary Figure S1B. **D)** Evolution of weight in mutant mice. Cross denotes death. Notably, starting at weeks 60-70, *Nek1^wt/t^ Pml^-/-^* mice, but not *Nek1^wt/t^*, exhibit significant weight loss of approximately 5%. Homozygous *Nek1^t/t^*mice show pronounced weight loss (16.4%), which is further exacerbated in the absence of *Pml*, reaching 20%. For clarity, *Pml^-/-^* controls are presented in Supplementary Figure S1B. **E)** Summary of the ALS-like symptoms. Evaluation of disease onset is described in Methods. **F)** Muscle strength and neuromuscular functions as measured by grip strength and hanging wire tests. In accordance with ethical and institutional regulations, no tests, analyses, or treatments involving animals could be conducted before they reached 6 weeks of age; this accounts for the gap observed in the graphs during the initial weeks. *Pml^-/-^*controls for grip strength, and trendlines, are presented in Supplementary Figure S1C. **G)** Western blot illustrating Nek1^t^ protein expression in spinal cords of animals with the indicated genotypes (*) wtNek1, (**) Nek1^t^. **H)** Immunofluorescence showing Nek1 protein in *wild-type* (top), *Nek1^wt/t^* (middle) or Nek1^t^ aggregates in *Nek1^t/t^* (bottom) spinal cord αMNs, identified by VAChT expression and size. Scale bars: 5 μm. Quantification of aggregates (30 αMNs from 3 mice for each genotype) is shown (right).

PML nuclear bodies (NBs) are membrane-less organelles, organized by the scaffold PML protein, regulating the activity or stability of client proteins through SUMOylation and ubiquitylation ^18–20^. *Pml^-/-^*mice exhibit no notable physiological symptoms, presenting a normal lifespan ^21,22^. Yet, PML may facilitate proteolysis of misfolded DRIPs (defective ribosomal products) or mutant proteins ^20,23^. Interferons boost NB-mediated SUMOylation by massively enhancing *PML* transcription, enhancing SUMO availability, and modulating oxidative stress ^20,24–26^. This drives recruitment of client proteins into these structures, subsequently promoting client *in situ* SUMOylation, and, for some of them, STUbL (SUMO-targeted ubiquitin ligase)- mediated ubiquitylation and proteasomal degradation ^20^.

In this study, we reveal that the truncated Nek1 protein (Nek1^t^) encoded by the *Nek1* Arg812Ter mutation is highly prone to aggregation in αMNs and drives ALS symptoms in mice. *Pml* is a key modifier of ALS penetrance, since *Pml* loss allows for ALS development even in heterozygotes (*Nek1^wt/t^*), while transcriptional *Pml* induction by poly(I:C) dramatically delays symptom onsets in homozygotes (*Nek1^t/t^*) by promoting Pml- and SUMO-dependent degradation of Nek1^t^, and crucially, clearing mutant protein aggregates, *in vivo*.

## RESULTS

### The ALS-linked Arg812Ter mutation yields a truncated Nek1 (Nek1^t^) protein in mice motoneurons

To explore NEK1^t^-associated ALS, we utilized CRISPR/Cas9 technology to modify mouse embryonic stem cells and introduced a point mutation into mouse *Nek1* gene, resulting in a premature stop codon mimicking the human Arg812Ter mutation (Figure 1B) ^13^. *Nek1^t^* heterozygous mice (*Nek1^wt/t^*) showed no significant phenotype (Figures 1C, 1D, 1E, 1F and Supplementary Figures S1A, S1C, S1D), in line with the incomplete ALS penetrance of *NEK1*-mutant human subjects ^4,13^. We demonstrated *Nek1^t^*expression in multiple mouse tissues, including spinal αMNs, using RT-PCR, Western blot and immunofluorescence (Figures 1G, 1H and Supplementary Figures S1E, S1F), demonstrating that the transcript is not subject to nonsense mediated RNA decay and that the pathogenic protein is expressed by the cells.

### ALS-linked NEK1^t^ is prone to aggregation

To clarify the molecular and cellular underpinnings of NEK1^t^-induced ALS, we expressed the GST-, GFP- or FLAG-tagged human mutant protein in HEK293 or Neuro2A cells. In all of these conditions, wild-type NEK1 protein (wtNEK1) exhibits a diffuse cytoplasmic expression (Figure 2A and Supplementary Figures S2A, S2B) ^6,7^. In contrast, the expression of tagged NEK1^t^ is confined to the nucleus, likely reflecting the loss of its two nuclear export sequences (Figures 1A, 2A and Supplementary Figures S2A, S2B) ^13^. Remarkably, NEK1^t^ forms aggregate-like nuclear or nucleolar structures, reminiscent of phase-separated Tau, α-synuclein, or TDP-43 aggregates (Figure 2A and Supplementary Figures S2A, S2B, S2C) ^27,28^. Nearly 80% of NEK1^t^-expressing cells present nuclear aggregates in either cell types, while none are observed in tagged NEK1-expressing cells (Figure 2A and Supplementary Figures S2A, S2B). Tracking the behavior of GFP-NEK1^t^ in real-time video-microscopy revealed that the mutant protein very rapidly formed nuclear aggregates (6 min), some of which later merged with each other, resulting in very large aggregates deposited within the nucleolus (Figure 2B). Fluorescence recovery after photobleaching (FRAP) analyses of GFP-NEK1^t^ aggregates showed poor fluorescence recovery after bleaching (Figure 2C), indicating a non-dynamic gel-like state ^27,28^. To biochemically evaluate the solubility of NEK1^t^, we employed sequential protein fractionation based on solubility in mild detergent (NP-40), followed by a strong denaturant (SDS). Wild-type NEK1 was primarily located in the NP-40-soluble fraction, indicating its soluble nature. In contrast, a very significant proportion of NEK1^t^ remained in the NP-40-insoluble, SDS-extractable fraction, revealing its aggregated/insoluble nature (Figure 2D), as observed with many pathogenic proteins associated to neurodegeneration. In nuclear aggregates of NEK1^t^, we observed substantial positivity for endogenous SUMO1 and SUMO2/3 by immunofluorescence, suggestive for post-translational modifications of the mutant protein (Figure 2E). We thus examined the SUMOylation of NEK1^t^ using two complementary methods — immunoprecipitation (Figure 3C, lanes 1-3) and proximity ligation assays (Supplementary Figure S2D) — which demonstrated that both wild-type NEK1 and its truncation mutant were conjugated by SUMO1 and SUMO2/3. Additionally, both proteins physically interacted with the SUMO-conjugating enzyme, UBC9, as demonstrated by proximity ligation assays (Supplementary Figure S2D). The NEK1^t^ mutant exhibited higher basal-level SUMO1 and SUMO2/3 interactions when compared to the wild-type protein (Supplementary Figure S2D). Importantly, SUMOylation was repeatedly linked to both enhanced solubility and degradation of aggregation-prone proteins ^19,29–33^.

**Figure 2:**
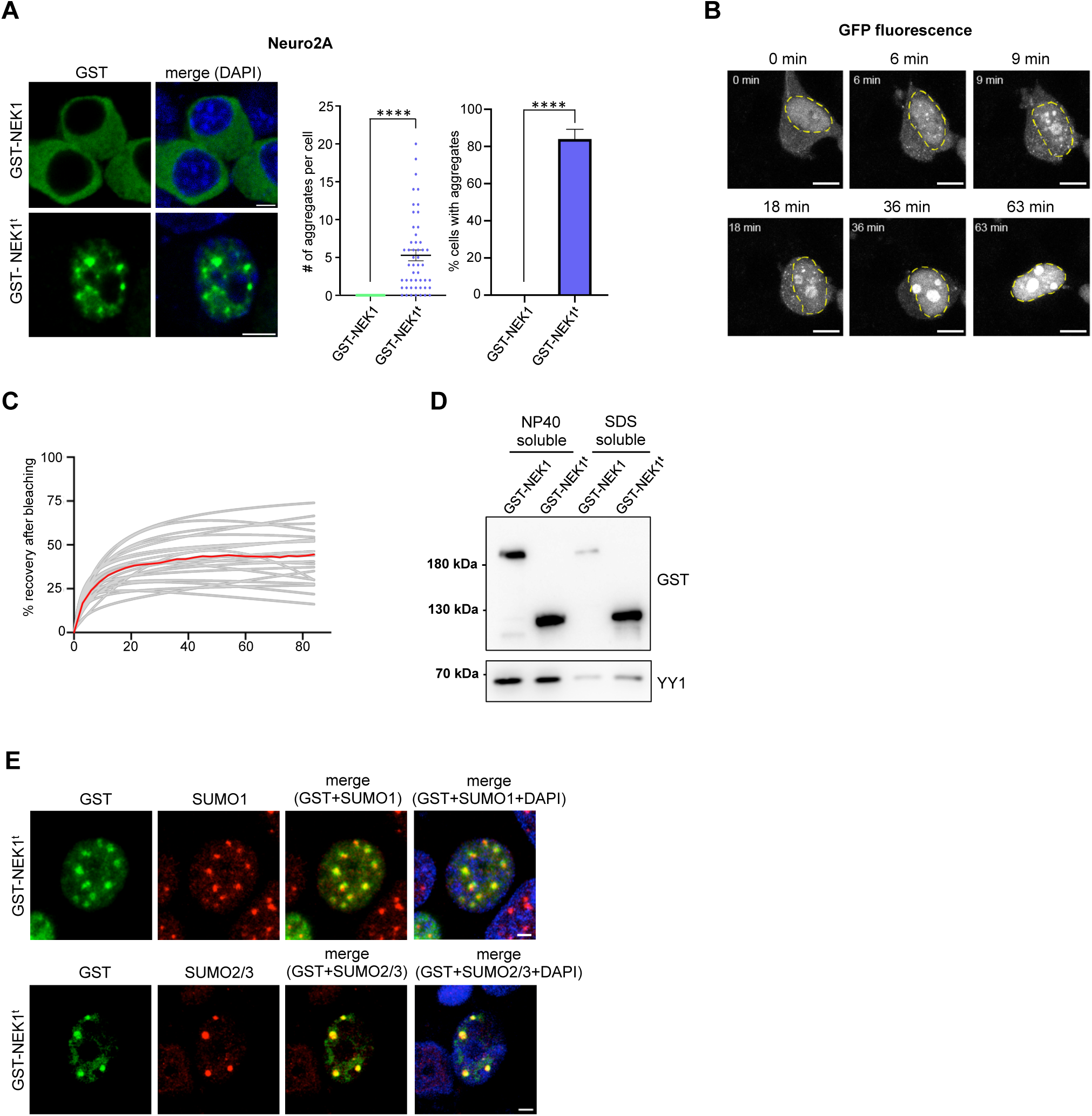
The insoluble and aggregation-prone nature of the truncated NEK1 mutant. **A)** Immunofluorescence illustrating NEK1^t^ protein aggregation in the nucleus (left). Quantification of these aggregates (right), n=3. Neuro2A cells were transfected with plasmids encoding GST-tagged NEK1^t^ or wtNEK1. Scale bars: 4.5 μm. **B)** Real time microscopy illustrates the dynamics of GFP-tagged NEK1^t^ aggregates in HEK293 cells. Dashed line: nucleus. Scale bars: 5 μm. **C)** FRAP analysis of NEK1^t^ protein aggregates**. D)** Assessment of NEK1^t^ protein solubility, in NP40, then SDS, in transiently transfected HEK293 cells. YY1, a nuclear transcription factor, was used as a positive control to confirm that the NP40-based lysis conditions employed were effective in extracting soluble nuclear proteins. **E)** In transiently transfected HEK293 cells, nuclear NEK1^t^ aggregates show strong positivity for endogenous SUMO1 (top) and SUMO2/3 (bottom) proteins. Scale bars: 2 μm.

**Figure 3:**
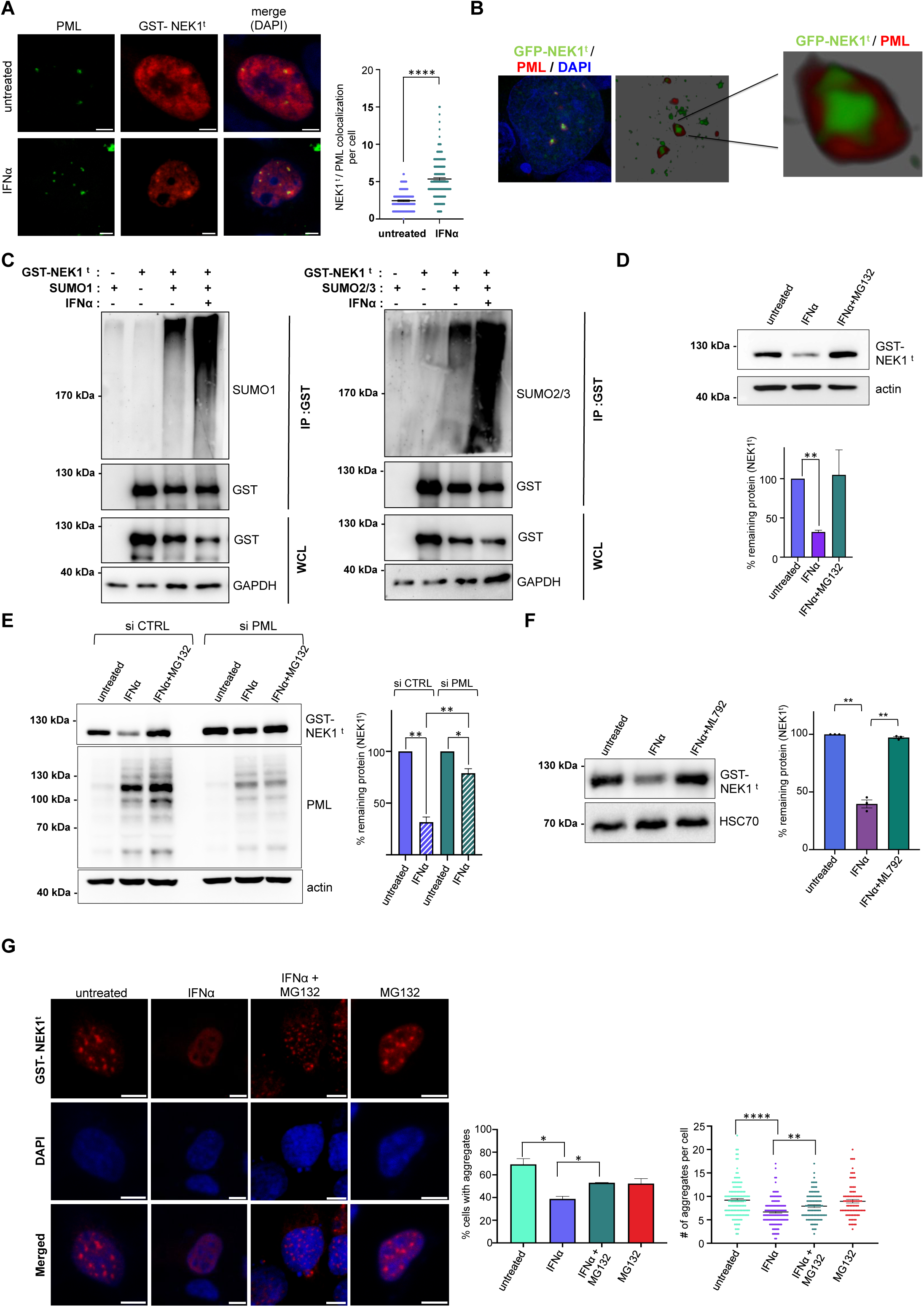
IFNα-induced PML NBs precipitate NEK1^t^ SUMOylation and degradation. **A)** A fraction of NEK1^t^ co-localizes with PML NBs in transfected HEK293 cells, a process inducible by exposure to IFNα. Quantifications are shown on the right (n=3). The untreated cell displays large NEK1^t^ aggregates. Scale bars: 4.5 μm. **B)** Recruitment of transfected GFP-NEK1^t^ into PML NBs (confocal image with deconvolution) following PML induction by IFNα. The full extent of the nucleus is also shown. **C)** NEK1^t^ modification by SUMO1 and SUMO2/3. HEK293 cells were transfected with the indicated expression vectors and treated or not with IFNα. IP: immunoprecipitation, WCL: whole cell lysate. **D)** IFNα treatment (24 hours) induces proteasome-dependent degradation of NEK1^t^ in HEK293 cells. MG132: proteasome inhibitor, n=3. **E)** *PML* silencing impedes IFNα-induced NEK1^t^ degradation. Quantifications (right), n=3. siPML: siRNA targeting *PML*, siCTRL: scrambled siRNA control. **F)** Inhibition of SUMOylation impedes IFNα-induced NEK1^t^ degradation. Quantifications (right), n=3. ML792: SUMOylation inhibitor. **G)** IFNα drives elimination of NEK1^t^ protein aggregates from HEK293 cells. Right, quantifications are shown (n=3). Scale bars: 8 μm.

### *Pml* mitigates ALS emergence through Pml-mediated, SUMO-facilitated degradation of Nek1^t^

Investigating potential genetic interactors of *Nek1^t^*, we focused on *Pml* gene, as a small fraction of NEK1^t^ co-localized with PML in nuclear bodies (NBs) in the basal state, while a very significant colocalization was observed following induction of PML by interferon alpha (IFNα) (Figures 3A, 3B). PML enhances partner conjugation by SUMOs, primarily SUMO2/3, which could enhance NEK1^t^ degradation and/or solubility to decrease aggregate formation ^19,20,34,35^. Strikingly, *Nek1^wt/t^* animals in a *Pml* knockout background (*Nek1^wt/t^ Pml^-/-^*) exhibited ALS-like phenotypes, characterized by severe muscle weakness, impaired coordination and motor function, starting 31.5 weeks post-birth or later, as measured by grip strength, hanging wire and limb clasping tests, with no such features observed in *Nek1^wt/t^* or *Pml^-/-^* controls (Figures 1E, 1F and Supplementary Figures S1C, S1D). They also developed significant weight loss (Figure 1D and Supplementary Figure S1B), all resulting in a markedly shortened lifespan (Figure 1C and Supplementary Figure S1B), though this phenotype was not fully penetrant. Both grip strength and limb clasping tests revealed an age-dependent, progressive decline in muscle strength and motor function in *Nek1^wt/t^ Pml^-/-^*mice (Figure 1F and Supplementary Figures S1C, S1D), underscoring the contribution of *Pml* loss to the motor deficits. Thus, *Pml* absence *in vivo* reveals ALS-like pathogenesis fueled by heterozygous Nek1^t^ expression.

In NEK1^t^-transfected cells, IFNα-induced recruitment of NEK1^t^ into PML NBs (Figures 3A, 3B) facilitated its hyper-conjugation by SUMO1 and SUMO2/3 (Figure 3C, lanes 4), as well as by ubiquitin (Supplementary Figure S3A), precipitating NEK1^t^’s proteasome-, PML- and SUMO-dependent degradation (Figures 3D, 3E, 3F and Supplementary Figure S3B). Indeed, IFNα−induced degradation was abrogated by the SUMOylation inhibitor ML972. Remarkably, this IFNα−enhanced, PML-dependent degradation process dramatically reduced the abundance of NEK1^t^ aggregates in transfected cells (Figure 3G). Such degradation of the mutant protein most likely explains the emergence of ALS-like symptoms in *Pml* null *Nek1^wt/t^* mice, where basal Nek1^t^ degradation is expected to be blunted.

In *Nek1^t^* homozygous mice (*Nek1^t/t^*), increased Nek1^t^ abundance and/or putative loss in Nek1 functions allowed for rapid development of ALS in *Pml* proficient background (median survival 35 weeks) (Figures 1C, 1D, 1E, 1F and Supplementary Figures S1C, S1D). Beyond ALS, *Nek1^t/t^* displayed ciliopathy-like phenotypes that can be attributed to the loss of some Nek1 functions, including dwarfism (Supplementary Figure S1A), polycystic kidneys, and male sterility (data not shown), since complete Nek1 ablation is embryonic lethal ^7–11^. Thus, the loss of the Nek1 C-terminus likely accounts for some of these features. *Nek1^t/t^* mice also present large numbers of Nek1^t^ aggregates within spinal cord αMNs, something never observed in wild-type or heterozygous animals (Figures 1H, 4B). *In vivo*, Nek1^t^ aggregates were observed in both nuclear and cytoplasmic compartments. A similar distribution was also observed in transfected cells following prolonged NEK1^t^ expression (data not shown), which may result from compromised nuclear membrane integrity or impaired nucleo-cytoplasmic trafficking. To explore any accelerating role of *Pml* loss, we generated *Nek1^t/t^* animals in a *Pml^-/-^* background. While these mice clearly displayed exacerbated muscle weakness, greater motor deficit and accelerated weight loss, they only succumbed a week earlier than *Pml* proficient *Nek1^t/t^* mice (Figures 1C, 1D, 1E, 1F and Supplementary Figures S1B, S1C, S1D) and exhibited a non-significant increase in Nek1^t^ aggregates in their αMNs (Supplementary Figure S1G). Overall, these findings demonstrate that Nek1^t^ drives an ALS-like phenotype, a process normally prevented by Pml, most likely through basal SUMOylation-induced Nek1^t^ degradation.

### *Pml* induction *in vivo* clears Nek1^t^ aggregates, alleviates disease symptoms and improves survival

We then assessed any potential therapeutic benefit of transcriptional induction of the *Pml* gene by IFNα on Nek1^t^-driven ALS-like phenotype *in vivo*. To induce the activation of interferon signaling *in vivo*, pre-symptomatic animals received intraperitoneal injections of poly(I:C) ^36–38^, twice a week, beginning at week 5 after birth, for the duration of their lifetime. Strikingly, treated *Nek1^t/t^* mice exhibited elimination of the Nek1^t^ proteins from multiple tissues within 3 weeks, including spinal cord (Figure 4A and Supplementary Figure S4A), and a significant reduction of Nek1^t^ aggregates in αMNs was observed in treated animals (Figure 4B). In contrast, wtNek1 expression levels were unaffected by poly(I:C) (Figure 4C and Supplementary Figure S4B). Functionally, poly(I:C)-treated *Nek1^t/t^*mice exhibited mild or no symptoms for several months and succumbed to ALS much later (median survival 54 weeks, compared to 35 weeks in untreated *Nek1^t/t^* mice) (Figures 4D, 4E and Supplementary Figure S4C). The clinical response to poly(I:C) therapy was observed in both sexes, indicating that the therapeutic effect was not sex-dependent (Supplementary Figure S4D). Critically, the same treatment regimen failed to mitigate any ALS-like symptoms or extend survival in *Nek1^t/t^ Pml^-/-^* animals (median survival 34 weeks) (Figures 4D, 4E and Supplementary Figure S4C) or to clear Nek1^t^ from tissues (Figure 4A and Supplementary Figure S4A). Overall, poly(I:C) therapy extended survival by 5 months, alleviating ALS-like symptoms, a process requiring *Pml* and associated to clearance of mutant protein aggregates in spinal cord αMNs.

**Figure 4:**
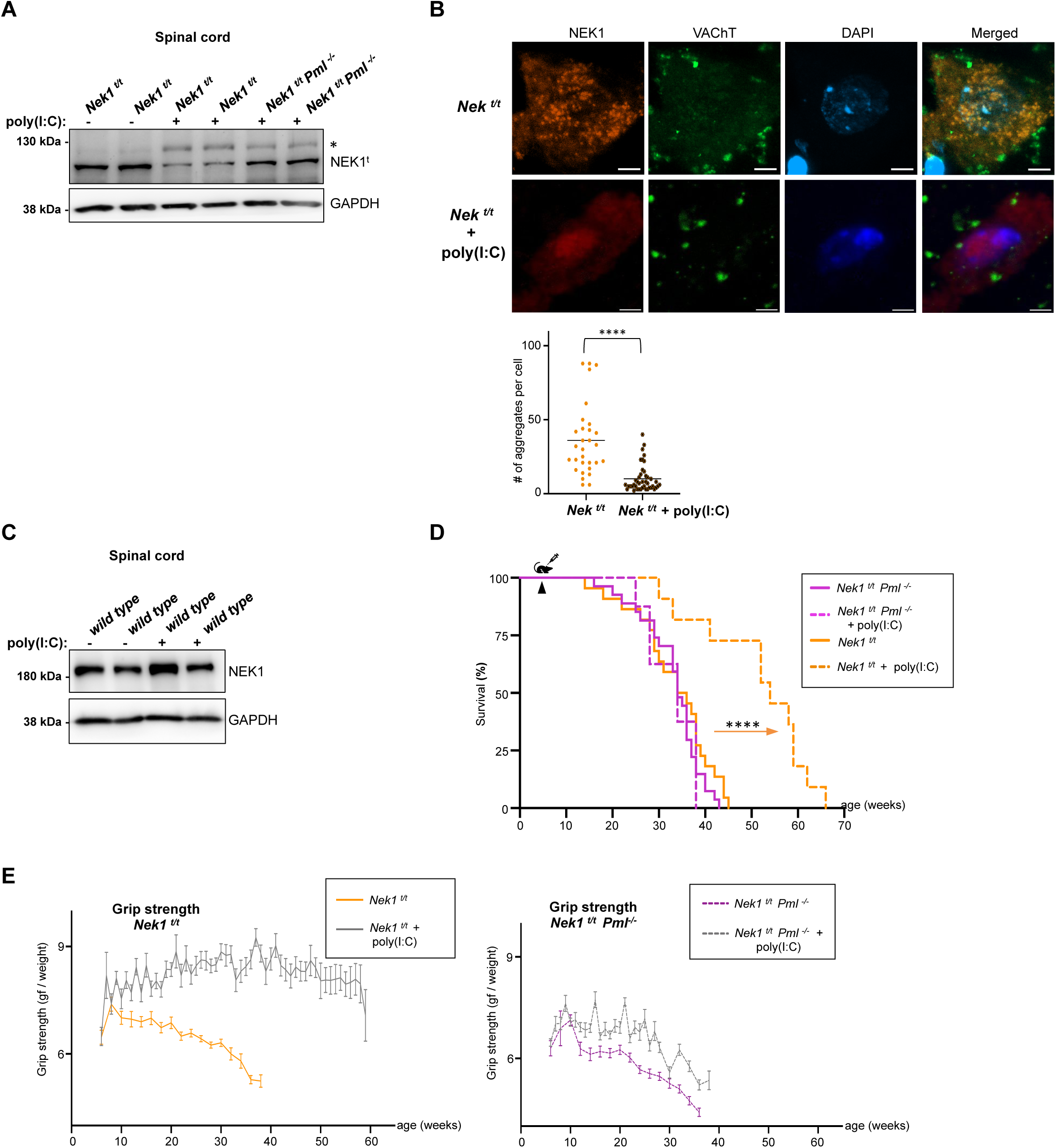
Therapeutic benefit of *Pml* induction by poly(I:C) through Nek1^t^ clearance *in vivo*. **A)** Western blot of Nek1^t^ protein from the spinal cord of age-matched *Nek1^t/t^* or *Nek1^t/t^ Pml^-/-^* mice following poly(I:C) therapy, (*) may correspond to SUMO-modified Nek1^t^. **B)** poly(I:C) therapy eliminates Nek1^t^ aggregates in αMNs of *Nek1^t/t^* mice. Quantifications: ≥30 αMNs from ≥3 mice. Scale bars: 5 μm. **C)** Western blot of wtNek1 protein from the spinal cord in *wild-type* mice following poly(I:C) therapy. Two different animals for each group. All animals in the study were of the same age. **D)** poly(I:C) therapy selectively extends lifespan in *Pml*-proficient *Nek1^t/t^* mice. Arrowhead: therapy initiation, continued until death. **E)** poly(I:C) therapy improves muscle strength and neuromuscular function selectively in *Pml*-proficient *Nek1^t/t^* ALS mice, as measured by grip strength. Trendlines in Supplementary Figure S4C. In accordance with ethical and institutional regulations, no tests, analyses, or treatments involving animals could be conducted before they reached 6 weeks of age; this accounts for the gap observed in the graphs during the initial weeks.

## DISCUSSION

Motoneuron degeneration in ALS is characterized by several pathological features, including dysregulated redox metabolism, impaired RNA transport, disrupted proteostasis, DNA damage accumulation or increased cellular excitability. Several ALS-linked mutations result in a toxic gain of function through protein aggregation, driving these cellular defects. Pharmacological targeting of the mutated aggregating protein may provide clinical benefits, as demonstrated by Tofersen, an FDA-approved antisense oligonucleotide, which blunts expression of mutated SOD1 proteins. While any specific role of nuclear NEK1^t^ aggregation in ALS remained uncertain, evidence presented here identifies a tight correlation between the expression of the truncated protein, the occurrence of aggregates in αMNs and ALS- like symptoms or disease progression. Moreover, *Pml* absence allows emergence of the disease in heterozygous *Nek1^wt/t^* mice, which likely reflects a role for Pml in SUMOylation and basal solubilization/degradation of the mutant protein mitigating its aggregation. Accordingly, eliminating Nek1^t^ aggregates from αMNs through poly(I:C)- induced *Pml* expression yields disappearance of ALS-like symptoms and survival benefits. In this setting, putative remaining defects of wild-type Nek1 function due to its homozygous C-terminal truncation may contribute to death ^12,16^. Very recently, PML was shown to promote solubilization of pathogenic TDP-43 variants in transfected cells ^35^. Our studies demonstrate that Nek1^t^ is degraded by the PML/SUMO/proteasome pathway, but this same pathway likely also promotes solubilization of the remaining mutant protein. The distinction between protein condensates and protein aggregates may be subtle, yet important, in the context of neurodegeneration. Several proteins implicated in neurodegenerative diseases, including ALS-associated factors (i.e. TDP-43, FUS), undergo phase separation to form biomolecular condensates. These condensates display dynamic, liquid-like properties and ability to fuse, features that we have observed here with truncated NEK1. However, under conditions of cellular stress, altered post-translational modification profiles or disease-associated mutations that promote misfolding, these dynamic condensates may transition into rigid, less dynamic and detergent-insoluble structures – hallmarks of pathological aggregation, as observed here with NEK1^t^, forming toxic, aggregate-like states. Overall, our observations shed a new light on the pathogenesis of NEK1-linked ALS, in line with the roles of protein aggregates in other neurodegenerative conditions. Yet, they highlight the key modulation of this process by the levels of PML and SUMO expression, supporting recent biochemical findings with another aggregating protein ^35^.

To our knowledge, this is the most dramatic benefit reported in treatment trials initiated pre-symptomatically in ALS mouse models. It remains to be seen whether this treatment will also be effective if initiated in symptomatic animals. While we demonstrate clear control of Pml abundance over Nek1^t^ aggregates, it is possible that other functions of Pml, notably metabolic ones ^39–42^, may also contribute to the beneficial outcome. *NEK1* mutations represent ALS-risk variants found in 2-5% of sporadic and familial cases, constituting a sizable proportion of ALS-related mutations ^3,4,12–14,16^. *NEK1* alterations may be found together with mutations in other susceptibility genes (i.e. *C9orf72, TARDBP*) that also yield toxic protein aggregates, which could reflect a mutual facilitation of phase separation ^4,14,43–45^. PML was proposed to facilitate solubilization and/or degradation of mutated ataxin-1, Huntingtin and TDP-43 proteins *in vitro* ^23,35^. Here we provide unambiguous demonstration of Pml-mediated degradation *in vivo*. In SCA1 mice, *Pml* absence somehow enhances pathological brain lesions, but not the clinical phenotype ^23^. Our study is the first to show, *in vivo*, a dramatic Pml-dependency of disease incidence, mutant protein stability and aggregation, as well as major Pml-dependent clinical response to interferon signaling activation and poly(I:C). Our observations clearly establish the *in vivo* feasibility of novel PML-based therapeutic strategies, providing support for therapeutic relevance of recent proposals that molecular glues bridging PML and misfolded partners can promote aggregate dissolution *in vitro* ^35^. Clearing disease-causing molecules through IFNα-induced PML NB formation could thus hold great promise as a treatment avenue for neurodegeneration, expanding the scope of PML targeting beyond its current applications in hematological cancers ^46,47^.

## METHODS

### Creation of a *Nek1^t^* ALS mouse model, genotyping, RNA and protein analyses

The mouse Nek1-Lys764Ter mutation corresponds to the human NEK1-Lys812Ter mutation found in a subset of ALS patients and results in the expression of the truncated Nek1^t^ protein. Nek1-Lys764Ter mutant mice and *Pml* deficient (*Pml^-/-^*) mice were created using the i-GONAD method ^48^. CRISPR/Cas9 reagents were injected into the oviducts of pregnant FVB/N females, followed by electroporation (NEPA21 Electroporator, Sonidel) of the entire oviduct. crRNA, tracrRNA, ssDNA, and Cas9 nuclease were purchased from IDT (Integrated DNA Technologies).

For Nek1-Lys764Ter mutation, the CRISPR target sequence (without PAM) was 5’- AACATACTCATACATACTTT-3’ and the ssODN sequence was 5’-GATGCAGTGACACTGGATACATCCTTCTCTGCAACCGAATGTATGTATGAGTATGTTAATGTCATGGCTTGGTTTATCAC-3’. For *Pml^-/-^* animals, the CRISPR target sequence (without PAM) was 5’- GTGTTGCACATGCGCGCTCC −3’.

Genotyping was performed by PCR and direct DNA sequencing for each animal. Earpieces were obtained from 3-week-old mice, after separation from their mothers. DNA extraction buffer containing proteinase K in a 1:100 ratio (20mg/ml, GoldBio, Cat # P-480-100) was added to each sample and left for overnight incubation in a shaking block (600rpm, at 58°C). The following day the samples were taken and following a series of washes with isopropanol and ethanol, genomic DNA was obtained and used for PCR reactions at the following conditions: Initial denaturation 95°C for 30 sec; [95°C for 30 sec, 60°C for 30 sec, 72°C for 45 sec]X34; final extension 68°C for 5 min. The following PCR primer sets were used:

*Nek1*-Forward 5’-ATACCCAGGAAGAAGAAATGGAAA-3;
*Nek1*-Reverse 5’- GCCACTGCAAAGACCAAACTT-3;
*Pml*-Forward 5’-GGTTGTCAGACTTGGCTGTGA-3;
*Pml*-Reverse 5’-GCAGCTGGACTTTCTGGTTCT-3’.

PCR with *Nek1*-Forward and *Nek1*-Reverse primers generates a 524bp fragment. The purified PCR product was then sequenced using Nek1-sequencing primer Fwd: 5’- GTGCTATTTCAGTAAGTACAG −3’. PCR with *Pml*-Forward and *Pml*-Reverse primers generates a 671bp fragment in wild-type animals and a 612bp fragment in the knockouts.

The expression of *Nek1* and *Nek1^t^* transcripts were verified by RT-PCR. RNA was isolated from from various mouse tissues using Direct-Zol RNA Isolation Kit (Zymogen, Cat # R2071). cDNA synthesis was performed using SensiFAST cDNA Synthesis Kit (Bioline, Cat # BIO-65053) according to manufacturer’s instructions. PCR was performed using primers specific for *Nek1* and *Gapdh* sequences. A set of primers was used to amplify a 167 bp-long shared region present in both wild-type and *Nek1^t^* transcripts. The following PCR primers sets were used:

*Nek1*-Forward 5’- TGGCAAACATGAAGCATCCA −3’;
*Nek1*-Reverse 5’- CACAAACCAGTCCAAAATCTGG −3’;
*Gapdh*-Forward 5’- AAGGGCTCATGACCACAG − 3’;
*Gapdh*-Reverse 5’ − CATCACGCCACAGCTTTCC −3’

(*Gapdh* was used as an internal control). The following PCR protocol was employed (initial denaturation 95°C for 30 sec; [95°C for 30 sec, 60°C for 20 sec, 72°C for 15 sec]X34; final extension 68°C for 5 min).

For protein and Western blot analysis, mouse spinal cord and heart tissues were removed and lysed in organic lysis buffer containing 50mM HEPES, 1% Triton X-100, 10mM EDTA, 2% SDS and 50mM NaCl. Protein lysates were mixed with 2X *L*aemmli buffer and boiled for 5 min at 70°C prior to SDS-PAGE. The following antibodies were used for the detection of proteins: human anti-NEK1 (Santa Cruz, Cat # sc-398813), human anti-GAPDH (Santa Cruz, Cat # sc-32233).

### Motor function tests

Grip strength tests (Ugo Basile, Cat # 47200) measure the maximum force that a mouse can exert on an apparatus, hence measuring muscle strength and overall neuromuscular function, documenting any possible muscle weakness. Each animal was placed on a base plate, being pulled by the tail. Three different measurements (of the force exerted, indicated on y-axes as gram force, gf) were taken for each animal, and the averages were calculated. The weekly scores of each animal were normalized to weight. Animals were subjected to the test starting at week 6 after birth.

For hanging wire tests, animals were allowed to hang on a wire and the time they were active and moving was recorded (indicated on y-axes as motility, units are in seconds, sec), within a timeframe of 4 minutes in total.

The limb clasping test is used to assess motor function impairments and to track disease progression in various rodent models of neurodegenerative disorders, in particular, in ALS. In this procedure, the animal is carefully lifted by its tail, and the posture of its hindlimbs is observed. Under normal conditions, healthy animals typically extend their hindlimbs outward, away from the body, when suspended. In contrast, animals with corticospinal or motor impairments exhibit an abnormal response, drawing their hindlimbs inward or "clasping" them toward the abdomen. The extent of this retraction is scored and used as a measure of motor dysfunction or neurological impairment. In Supplementary Figure S1D, the Y-axis represent the severity of motor function impairment, where 3 indicates the most severe impairment and 0 corresponds to normal motor function.

The onset of ALS symptoms was evaluated through multiple factors, including decline in muscle strength and overall neuromuscular function (measured by grip strength and hanging wire tests), motor deficits and coordination loss (evaluated through limb clasping and rotarod tests, with the latter data not shown), and weight loss. A 20% reduction in muscle strength was considered the onset of symptoms.

### *In vivo* treatments

In order to induce the activation of endogenous interferon signaling and *Pml* gene expression *in vivo* ^36–38^, the animals received intraperitoneal injections of poly(I:C) (Invivogen, Cat # 31852-29-6) twice a week, beginning at week 5 after birth, at a dosage of 20μg per gram. Treatment was continued over the lifespan of animals (or until sacrifice).

### Immunofluorescence on tissues

Animals were sacrificed in a CO_2_ chamber and the spinal cords were obtained after dissection, followed by fixation in 4% paraformaldehyde for an hour at 4°C, then overnight incubation in 20% sucrose for better cryoprotection. Tissues were subsequently embedded in molds filled with OCT (embedding matrix, VWR, Cat # 65608-930) and frozen. Tissue sections were obtained in 30μM thickness using Cryostar NX 70 cryostat. Immunofluorescence was performed on tissue sections following cytoplasmic and nuclear permeabilization of this highly myelinated tissue. For this, after washing away the OCT using PBS, slides were placed in 2% Triton X-100 in PBS for 30 min at room temperature to remove the myelin, followed by the “heat shock demasking” step, performed by incubation in citrate buffer (0.1M citric acid and 0.1M sodium citrate, pH 6.0) for 20 min at 90°C. Following subsequent incubation in PBS containing 3% H_2_O_2_ and 10% methanol for 15 min at room temperature, the tissues were permeabilized in 0.5% Triton X-100 (1 hr at room temperature). Next, slides were washed in PBS and then incubated in 100mM glycine, washed again in PBS and finally placed in a blocking buffer containing 1% Fetal Bovine Serum (Gibco, Cat # 10270106) and 1% Triton X-100. Primary antibody incubation was performed overnight +4°C.The next day slides were washed three times in PBS containing 0.1% Triton X-100 and incubated with secondary antibodies for 2 hrs at room temperature. After washing three times in PBS, slides were mounted in DAPI (Southern Biotech, Cat # 0100-20). The following antibodies were used: human anti-NEK1 antibody (Santa Cruz, Cat # sc-398813) and anti-VAChT (Sigma-Aldrich, Cat # ABN100). Images were obtained using a confocal microscope (Leica TCS SP8, USA) and analyzed using Image J, Fiji and LASX software packages. Alpha-motoneurons were detected based on size and VAChT expression.

### Cell culture, constructs, transfections and treatments

HEK293 or Neuro2A cells were maintained in DMEM supplemented with 10% FBS and 1% penicillin/streptomycin. Cells were kept in a humidified incubator that maintained the temperature at 37°C and CO_2_ levels at 5%. The expression plasmid for the GST-tagged human NEK1 was purchased from Addgene. In order to generate the NEK1^t^ mutant, which contains a premature stop codon at amino acid position 812, site directed mutagenesis was performed on the abovementioned plasmid using the following primers:

Arg812Ter-Forward: 5’-CTACAACTGAATGACATACAGTG-3’;
Arg812Ter-Reverse: 5’-CACTGTATGTCATTCAGTTGTAG-3’

Plasmids encoding SUMO and ubiquitin proteins were described previously ^20,49^. The day before plasmid DNA transfection, HEK293 cells were seeded as they reach 50-70% confluency at the time of transfection. 500ng or 1μg plasmid DNA were transfected into cells grown in 12- and 6-well plates, respectively. For transfection to 12-well plates, the plasmid DNA was dissolved in water. Then 30.5μl of 2M ice-cold CaCl_2_ was added to the mixture, followed by dropwise addition of 250μl 2x HBS (Lonza, Cat # CC-5024). After 10 min incubation, transfection mix was gently added to the cells.

siRNA transfections of HEK293 cells were performed using HiPerFect transfection reagent (Qiagen, Cat # 301705), according to manufacturer’s instructions. The siRNA sequence used to silence Pml expression was 5’- AAGAGTCGGCCGACTTCTGGT-3’. The control (non-targeting) siRNA sequence used in the same experiment was 5’-UUCUCCGAACGUGUCACGUTT-3’.

Neuro2A cells were transfected using the Metafectene reagent (Biontex, Cat # T020-1.0), according to manufacturer’s instructions.

HEK293 cells were treated with recombinant human interferon alpha (PBL Assay Science, Cat # 11100-1) for 24 hours at a concentration of 1000 U/ml. MG132 (Hycultec, Cat # HY-13259) treatment was performed at a final concentration of 1μM (24 hours). ML792 (MedKoo Biosciences) treatment was also performed at a final concentration of 1μM (24 hours). Neuro2A cells were treated with recombinant mouse interferon alpha (Sino Biological, Cat # 50525-M01H) for 24 hours at a concentration of 1000 U/ml.

The following antibodies were used for Western blots in HEK293 or Neuro2A cells: anti-GST (Cell signaling, Cat # 2624S), human anti-GAPDH (Santa Cruz, Cat # sc-32233;), human anti-actin (BioLegend, Cat # 622102), human anti-YY1 (Santa Cruz, H-10, Cat # sc-7341), human anti-HSC70 (Santa Cruz, Cat # sc-7298), human anti-PML (Santa Cruz, H-238, Cat # sc-5621 and Santa Cruz, E-11, Cat # sc-377390), human anti-SUMO1 (CST, Cat # 4930), human anti-SUMO2/3 (Abcam, Cat # ab3742).

### Immunoprecipitations and SUMOylation assays

SUMOylation and ubiquitylation assays on NEK1 or NEK1^t^ proteins using immunoprecipitation were performed as described previously ^20,49,50^, following transfection of HEK293 cells with plasmids encoding the GST-tagged versions of these proteins and/or SUMO1, SUMO2/3 or ubiquitin proteins. Cells were treated with recombinant human interferon alpha (PBL Assay Science, Cat # 11100-1) for 24 hours at a concentration of 1000 IU/ml. An anti-GST antibody (Cell Signaling, Cat # 2624S) was used for immunoprecipitation. PLA-based SUMOylation assays (Duolink, Sigma Aldrich) probing interactions with endogenous SUMO1 or SUMO2/3 proteins or with the E2 SUMO-conjugating enzyme UBC9 were performed as described previously ^49^.

### Solubility Assay

In order to assess the solubilities of wild-type and NEK1^t^ proteins, plasmids encoding these proteins were expressed in HEK293 cells. Initially, we obtained a fraction of NP-40-soluble proteins, followed by centrifugation to separate NP-40-insoluble proteins, which were subsequently resolubilized with SDS, representing aggregates. Cells were collected 48 hrs post-transfection and lysed in 150 μl NP40-containing buffer (50mM Tris-HCl pH 8.0, 150mM NaCl, 1% NP40) by incubation for 30 min on ice. Next, the supernatant was collected after centrifuging the samples at 13000 rpm for 20 minutes at 4°C (this is the NP40-soluble fraction) and mixed with 50μl 4x Laemmli buffer and boiled. The remaining pellet was lysed by boiling in 10 μl of 2x Laemmli buffer (this is the SDS-soluble fraction). Samples were subsequently analyzed by Western blotting. YY1 is a nuclear protein (transcription factor) and was used as a positive control to demonstrate that the NP40-based lysis conditions that we used allowed extraction of soluble nuclear proteins.

### Immunofluorescence, confocal microscopy, PLA, FRAP and live cell imaging

The following antibodies were used for immunofluorescence on HEK293 or Neuro2A cells: human anti-GST (Cell Signaling, Cat # 2624S), human anti-FLAG (Sigma, Cat # F1804). Coverslips were placed in 12-mm culture dishes where HEK293 or Neuro2A cells were seeded. Cells were fixed in 4% PFA solution (20 min at room temperature). Following 2 rinses in cold PBS, cells were permeabilized in 0.25% Triton X-100 (15 min). They were then washed 3 times with PBS (5 min each) and blocked in 1% BSA (in PBS-Tween) for 1 hr. Primary antibody incubation was performed in blocking solution for 1 hr at room temperature in a humidified chamber, followed by 3 washes in PBS-Tween (5 min each). Secondary antibody incubation was performed in blocking solution for 1 hr at room temperature, this time in dark, followed by 3 washes in PBS-Tween (5 min each). Next, cells were incubated in DAPI (1 μg/ml, Southern Biotech, Cat # 0100-20) for 3 min, which was followed by rinsing in PBS. Coverslips were mounted on slides using the fluoroshield mounting medium. Images were acquired using a confocal microscope (Leica TCS SP8, USA) and analyzed using Image J, Fiji and LASX software packages.

For real-time microscopy analyses, a Leica SP8 confocal microscope with an incubation chamber was used. Unsynchronized cells were analyzed at 37°C in a 5% CO_2_ atmosphere at 0.375-second intervals in a 512×512 pixel format using an HC PL APO CS2 40x 1.3 NA objective.

For deconvoluted confocal analysis, high-resolution images were captured using an HC PL APO 100x/1.4 NA CS2 objective and deconvoluted with Huygens software.

For fluorescence recovery after photobleaching (FRAP) analyses, the FRAP module on the Leica SP8 microscope was used. Bleaching (fluorescence signal elimination) was performed with a 488 Argon laser at 100% power capacity. Images were captured using Leica LASX software and analyzed with Image J.

PLA (Proximity ligation assay, Duolink; Sigma-Aldrich) was performed as previously described ^20,50,51^.

### Statistical analyses

For Figure 1C, animals from five different genotypes were monitored for a thorough lifespan comparison. The groups included 12 *wild-type*, 14 *Nek1^wt/t^*, 22 *Nek1^t/t^*, 18 *Nek1^wt/t^ Pml^-/-^* and 27 *Nek1^t/t^ Pml^-/-^* animals. Statistical analysis was performed using the Log-Rank (Mantel-Cox) and Gehan Generalized Wilcoxon Tests. For Figure 1D, the weights of animals from five different genotypes were monitored up to week 70 after birth (or until their week of death). For each group, ≥7 animals were included in the analysis. Statistical analysis was performed using the Student’s t-test (paired).

ALS-associated weight loss in *Nek1^t/t^* mice (16.4%) was accelerated in *Pml* absence (20%), as calculated by animals’ weight at week of death (indicated by a cross) versus maximal weight attained (week 25). *Nek1^wt/t^ Pml^-/-^* mice, but not *Nek1^wt/t^*, also exhibit weight loss starting at weeks 60/70. In Figures 1F and 4E, each data point corresponds to the average of measurements from at least 5 different animals for a given week (and at least 4 different animals for each group in Supplementary Figure S1D), with a minimum of three animals used at very late time points as some animals begin to die. In Supplementary Figures S1B and S1C, ≥4 *Pml^-/-^* animals were monitored. In Figure 4D, animals from two different genotypes that received therapy or not were monitored for survival comparison. The groups included 22 *Nek^t/t^*, 27 *Nek1^t/t^ Pml^-/-^*, 11 *Nek^t/t^* receiving poly(I:C) and 8 *Nek1^t/t^ Pml^-/-^* receiving poly(I:C). In Supplementary Figure S4D, number of animals for each group was as follows: 7 untreated males, 6 treated males, 7 untreated females, 5 treated females. Statistical analysis was performed using the Log-Rank (Mantel-Cox) and Gehan Generalized Wilcoxon Tests. For the quantifications of aggregates in motor neurons and transfected cells, as well as of protein abundances in Western blots, and also of positive signals in PLA experiments, statistical analyses were performed using the Student’s t-test (unpaired, assuming unequal variances). The asterisks indicate the following p-values: (*) <0.05, (**) <0.01, (***) <0.001, (****) <0.0001.

## Supporting information

Revised supplementary figures

## ACKNOWLEDGEMENTS

This work was supported by an EMBO Installation Grant (IG3336 to U.S.) from the EMBO Young Investigator Network (YIN) Program, a TÜBİTAK–France Bilateral Grant (No. 119N095 to U.S. and H.d.T.), and an EMBO Scientific Exchange Grant (to U.S. and A.P.)

## ABBREVIATIONS

ALS: amyotrophic lateral sclerosis
DRIPs: defective ribosomal products
GFP: green fluorescent protein
IFNα: interferon alpha
αMN: alpha-motoneuron
NB: nuclear body
NEK1: NIMA-related kinase 1 (human protein)
NEK1^t^: truncated NEK1 protein (human)
Nek1: NIMA-related kinase 1 (mouse protein)
Nek1^t^: truncated Nek1 protein (mouse)
PML: promyelocytic leukemia (human protein)
Pml: promyelocytic leukemia (mouse protein)
poly(I:C): polyinosinic:polycytidylic acid
SCA1: spinocerebellar ataxia type 1
SCA7: spinocerebellar ataxia type 7
SUMO: small ubiquitin-like modifier
STUbL: SUMO-targeted ubiquitin ligase
TDP43: TAR DNA-binding protein 43
wtNEK1: wild-type NEK1 protein

## SUPPLEMENTARY FIGURE LEGENDS

**Supplementary Figure 1:** Expression of a truncated, aggregation-prone Nek1 mutant (Nek1^t^) induces an ALS-like phenotype in mice. **A)** *Nek1^t/t^* mice display multiple symptoms including dwarfism. **B)** Survival of the *Pml^-/-^*control group is comparable to that of *wild-type* mice, consistent with previous studies and established findings in the literature. At the time of submission, *Pml^-/-^* mice were 50 weeks old, remained healthy, and continued to be monitored (left). Evolution of body weight in the *Pml^-/-^* control group up to week 50, compared to *wild-type* mice (right). **C)** Trendlines for the grip strength and hanging wire analyses shown in Figure 1F (top). Muscle strength and neuromuscular functions in *Pml^-/-^*control mice remained normal and showed no decline compared to *wild-type* and *Nek1^wt/t^* mice, as assessed up to week 50 (bottom, and also see Supplementary Figure S1D). **D)** Motor function in mice, categorized into three age groups as indicated, was assessed using the limb clasping test. The Y-axis represents the severity of motor function impairment, where 3 indicates the most severe impairment and 0 corresponds to normal motor function. *Pml^-/-^*controls are also shown. **E)** RT-PCR from mouse tissues demonstrates the expression of *Nek1^t^* transcripts in animals of the indicated genotypes. A set of primers was used to amplify a 167 bp-long shared region present in both *wild-type Nek1* and *Nek1^t^* transcripts. NC: negative control. **F)** Western blot illustrating the expression of Nek1^t^ protein in the heart tissue of animals with the indicated genotypes (*) wtNek1, (**) Nek1^t^. **G)** Immunofluorescence showing Nek1^t^ protein aggregates in spinal cord αMNs (identified by size and VAChT expression) of *Nek1^t/t^ Pml^-/-^* mice (bottom panel), compared to *Pml*-proficient *Nek1^t/t^*animals (top panel). Quantification (right) shows the number of aggregates (30 αMNs from 3 different mice for each genotype). Scale bars: 5 μm.

**Supplementary Figure 2:** Further characterization of NEK1^t^ and its aggregates. **A)** Immunofluorescence illustrating NEK1^t^ protein aggregation in the nucleus (left). Quantification of these aggregates (right), n=3. HEK293 cells were transfected with plasmids encoding GST-tagged NEK1^t^ or wtNEK1. Scale bars: 11 μm. **B)** Immunofluorescence illustrating NEK1^t^ protein aggregation in the nucleus (left). Quantification of these aggregates (right), n=3. HEK293 cells were transfected with plasmids encoding GFP-tagged or FLAG-tagged NEK1^t^ or wtNEK1. Scale bars: 5 μm. **C)** Immunofluorescence reveals NEK1^t^ protein aggregates within the nucleolus. Fibrillarin serves a nucleolar marker. Scale bar: 10 μm. **D)** Proximity ligation (Duolink) assays (PLA) were used to investigate the physical interaction of wtNEK1 or NEK1^t^ with endogenous SUMO1 and SUMO2/3 peptides, and the UBC9 E2 SUMO conjugating enzyme, suggesting post-translational modification of these proteins with SUMO in HEK293 cells (transfected with GST-tagged proteins). Z-stack projections are shown. Nuclei were stained with DAPI (scale bars: 5 μm). Notably, the PLA interaction signals were predominantly cytoplasmic for wtNEK1, whereas they were primarily nuclear for NEK1^t^, consistent with the respective localization patterns of these proteins. Quantifications of positive Duolink signals are shown in the graph. n>30 cells per experiment, data represent mean values from two experiments per condition, including the negative controls using a single antibody of a given Duolink pair, ± SEM (****P < 0.0001, unpaired t-test).

**Supplementary Figure 3:** IFNα-induced NEK1^t^ ubiquitylation and degradation. **A)** NEK1^t^ is heavily ubiquitylated following IFNα treatment. HEK293 cells were transfected with the indicated expression vectors and treated or not with IFNα. IP: immunoprecipitation, WCL: whole cell lysate (where HSC70 serves as a loading control). SUMO2/3 was co-expressed to facilitate potential SUMO-dependent ubiquitylation of NEK1^t^, which is then strongly induced by IFNα. **B)** IFNα treatment (24 hours) induces proteasome-dependent degradation of transfected GST-NEK1^t^ in Neuro2A cells. MG132: proteasome inhibitor.

**Supplementary Figure 4:** *Pml*-dependent Nek1^t^ degradation and clinical response to poly(I:C) therapy. **A/B)** Western blot of Nek1^t^ protein from the heart tissue of *Nek1^t/t^* or *Nek1^t/t^ Pml^-/-^*mice (**A)** or of wtNek1 protein in *wild-type* mice (**B)**, following poly(I:C) therapy. Two different animals for each group. All animals in the study were of the same age. **C)** Trendlines for the grip strength analyses shown in Figure 4E. **D)** A robust clinical response to poly(I:C) therapy was observed in both male and female mice.

