## Supplementary material for "*Pml* loss worsens NEK1-linked ALS and *Pml* induction drives NEK1 degradation, precluding disease onset": Revised supplementary figures

### Supplementary Figure 1

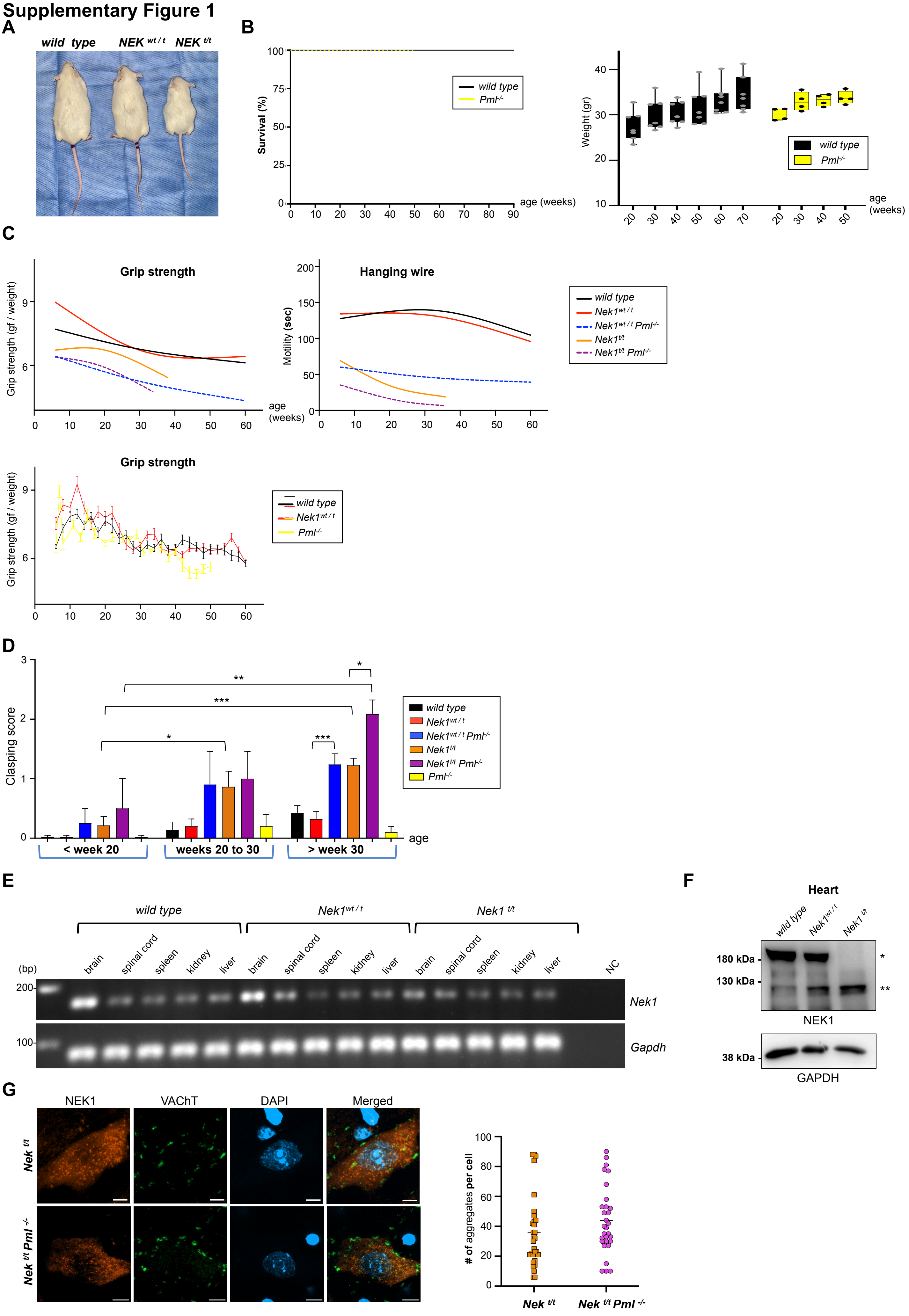

### Supplementary Figure 2

A

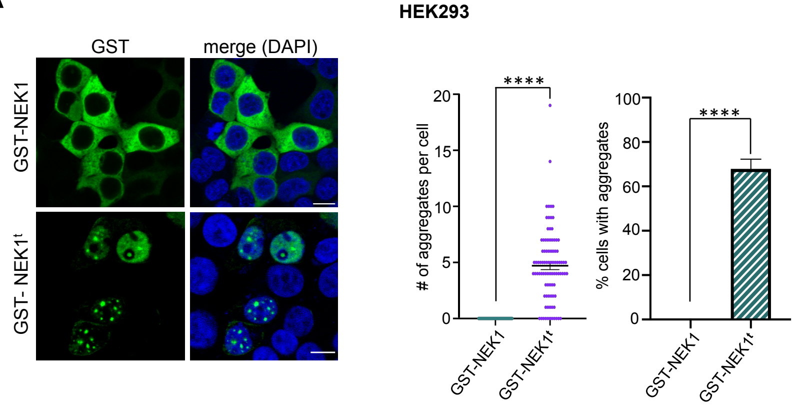


B

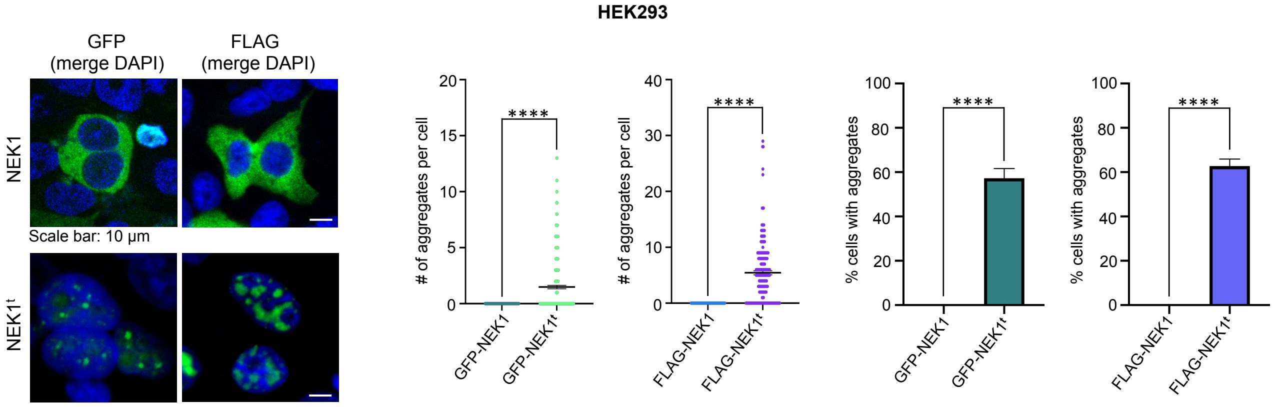


C

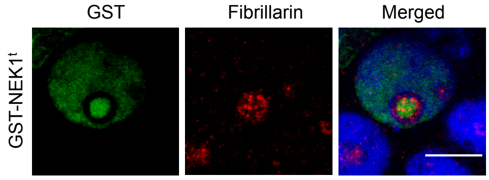


D

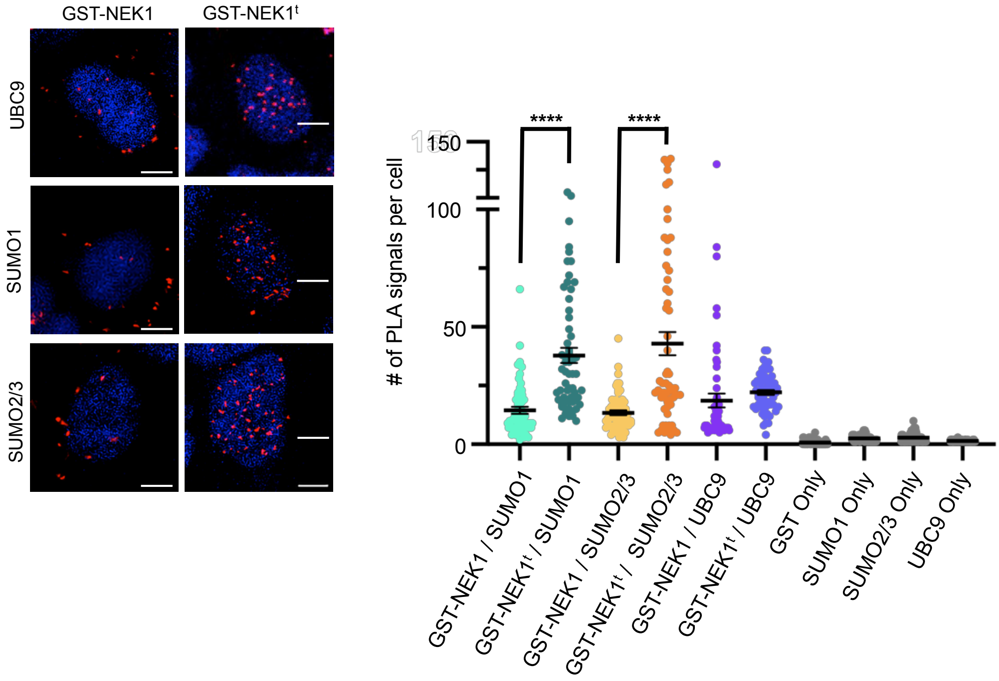

### Supplementary Figure 3

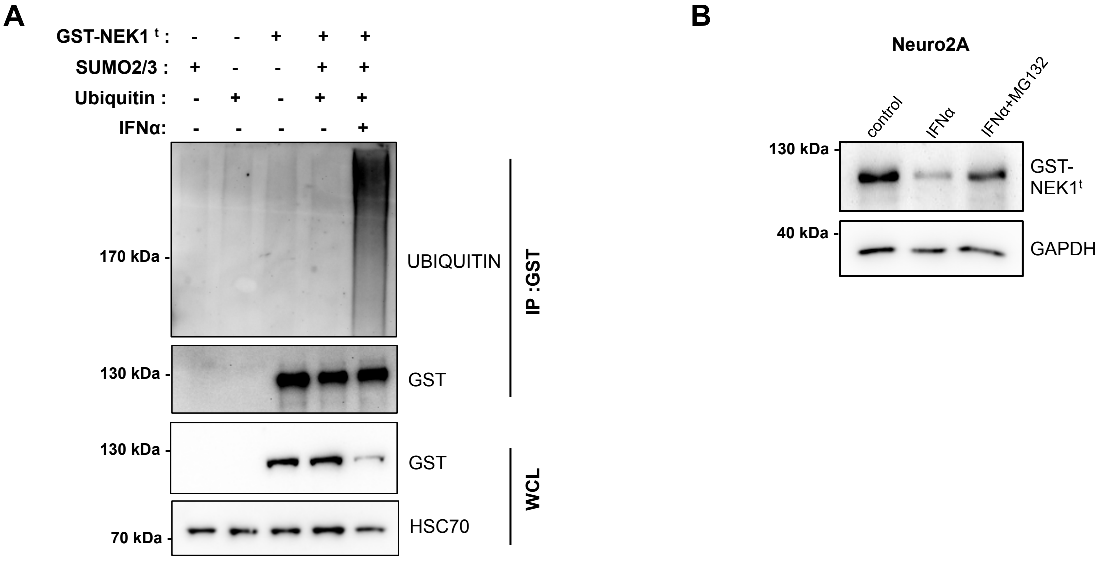

### Supplementary Figure 4

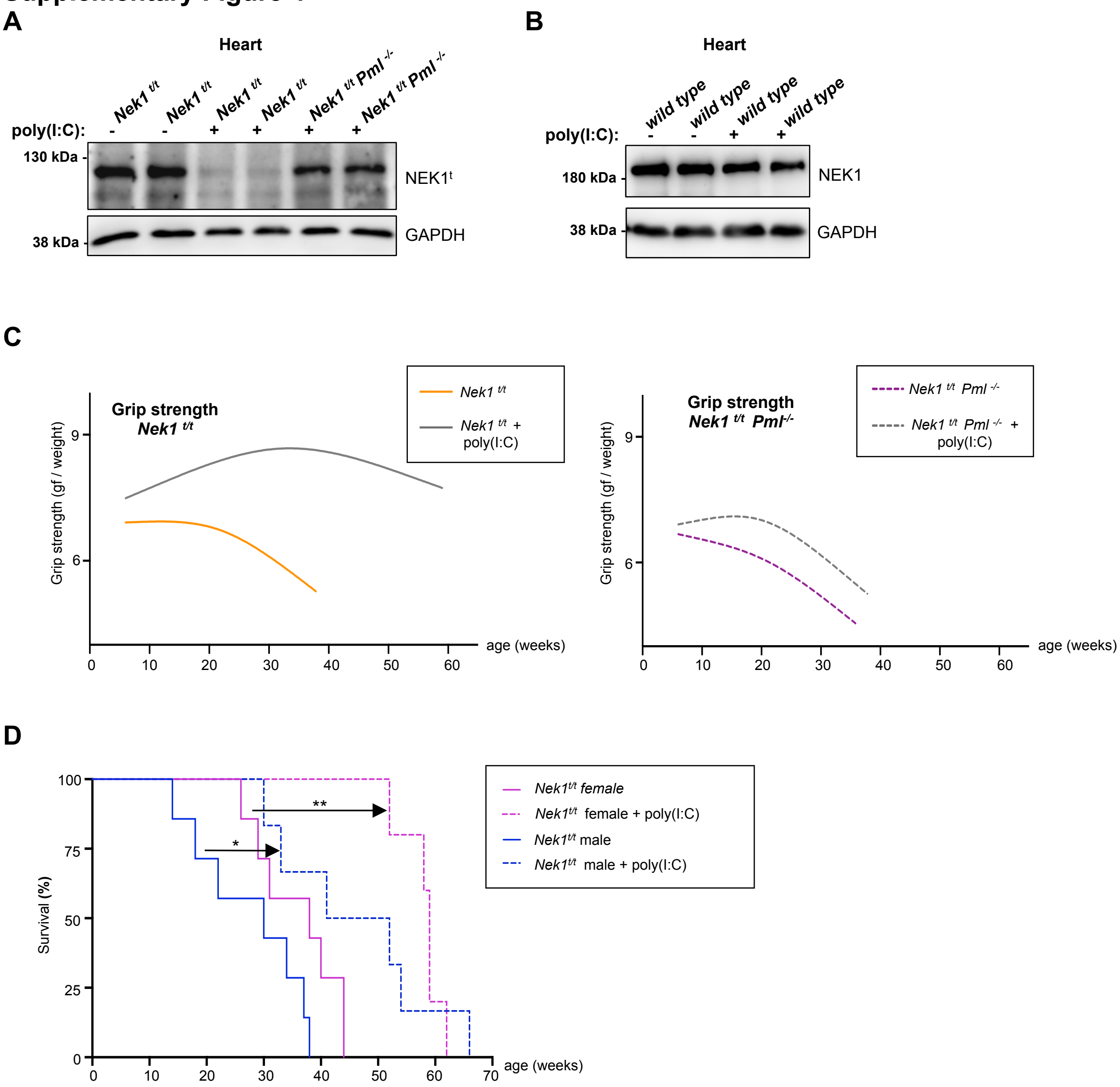
